# *ttc39bl,* one of the two duplicated paralogs of the *tetratricopeptide repeat domain 39B* gene, is essential for carotenoid coloration in medaka (Oryzias latipes) and is the gene responsible for the *r* locus

**DOI:** 10.1101/2025.06.15.659736

**Authors:** Tetsuaki Kimura, Ituro Inoue

## Abstract

Body color plays key roles in fitness, communication with others, and hiding. In poikilothermal vertebrates, the body color is mainly determined by types and distributions of chromatophores. Among them, carotenoid color of xanthophores/erythrophores is important for interspecific diversity and colorful males as sexual dimorphism. Most vertebrates cannot synthesize carotenoids in their bodies and must ingest them from food. However, the genes involved in the uptake process are not fully understood. Therefore, we tried to identify the causal gene of the carotenoid color mutant of medaka. The HdrR-II1 strain used in the genome project has orange body color in males and white body color in females. The orange and white body color was known to be controlled by the sex-linked *R* locus, but the causal gene of this was unknown. In this study, we identified that the causal gene of the *R* locus is *tetratricopeptide repeat domain 39b like* (*ttc39bl*). In the HdrR-II1, the *ttc39bl* on the Y chromosome is normal, but the *ttc39bl* on the X chromosome has an 821 bases insertion in exon 3 and is broken. This insertion is also present on both the X and Y chromosomes of commercially available white medaka.

## Introduction

In vertebrates, body color is known to be a critical factor for hiding and communicating with each other, thereby it has substantial effects on fitness. Body color mainly depends on types and distributions of chromatophores, which are derived from the neural crest in poikilothermal vertebrates (Kelsh et al., 2009; Sauka-Spengler and Bronner-Fraser 2008). Most chromatophores have pigments that determine their color: Melanocytes have black pigment, xanthophores have yellow pigment, and erythrophores have red pigment (Schartl et al., 2016). Meanwhile, iridophores have light-reflecting substances that enable them to achieve iridescent colors.

Carotenoids are one of such pigment, used in xanthophores/erythrophores. Carotenoids colors provide multiple roles in interspecific diversity, sexual dimorphism, and nuptial colors (Blount and McGraw 2008). Most vertebrates cannot synthesize the carotenoids in their bodies, thus need to intake them from food. In intestine, food-derived carotenoids are incorporated with other lipophilic substances into mixed micelles composed of bile acids, phosphate, cholesterol, etc., and taken up into the enterocyte by membrane transporters such as S*cavenger receptor class B member 1* (SCARB1), CD36, or *NPC1 Like Intracellular Cholesterol Transporter 1* (NPC1L1). In enterocyte, carotenoids are packaged into chylomicrons by the apoB-dependent pathway, secreted from the basolateral membrane, and ultimately taken up by the body (Reboul 2019). Some carotenoids are taken up by the non-apoB-dependent pathway mediated by the ABCA1 transporter (Reboul 2019). Carotenoids are concentrated and stored as oil droplets in specialized organelles within xanthophores/erythrophores (Granneman et al., 2017).

So far, several genes are known to be involved in the carotenoid coloration in chromatophores. S*CARB1* is a high-density lipoprotein receptor and essential for carotenoid accumulation in vertebrates (Toomey et al., 2017; Saunders et al., 2019). *perilipin 6* (*Plin6*) is critical for lipid droplet formation and acts carotenoid accumulation in teleost xanthophores (Granneman et al., 2017). *Tetratricopeptide Repeat Domain 39B* (*TTC39B*) is involved in lipid metabolism (Teslovich et al., 2010; Hsieh et al., 2016) and carotenoid coloration in vertebrates (Hooper et al., 2019; Salis et al., 2019; Ahi et al., 2020). In addition, *Beta-carotene oxygenase 2* (*BCO2*) and *CYP2J19*, members of the cytochrome P450 family of monooxygenases, are known to be involved in carotenoid coloration in vertebrates (Eriksson et al., 2008; Toews et al., 2017; Twyman et al., 2016; Mundy et al., 2016). However, the interactions between these genes are not known. So, the current information is insufficient for full understanding of how erythrophores and xanthophores colorations are achieved. Therefore, we intended to identify the responsible gene of the *r* locus involved in xanthophore pigmentation in medaka (*Oryzias latipes*).

In medaka, there is white medaka as a deficient of xanthophores pigmentation. White medaka has been maintained as a pet since Edo period, about 190 years ago in Japan (Ishikawa et al., 2022). Aida reported that sex-linked *r* locus is responsible for white body color (Aida 1921). Medaka has a male heterogametic sex-determining system, i.e. XY for male and XX for female. The sex-determining gene was identified as *dmy* locating on chromosome 1 corresponding to functional Y chromosome. The homologous chromosome lacking *dmy* corresponds to functional X chromosome (Matsuda et al., 2002, 2007). An inbred strain of medaka, HdrR-II1, has the recessive locus of *r* on the X chromosome and the dominant locus of *R* on Y chromosome (Hyodo-Taguchi 1996). As the result, HdrR-II1 female has white body color and male has orange body color (Fig. 1). It is known that medaka homozygous for the *r* locus have xanthophore without carotenoid colored in the adult stage (Hama 1975). The *r* locus locates 0.36 cM apart from the *dmy* and 0.18 cM apart from a genetic marker, 51H.7F, on chromosome 1 according to the linkage analyses (Matsuda et al., 2002). 51H.7F locates in *ATP-binding cassette sub-family G member 2d* (*abcg2d*) (Fig.S1), however, the physical location of *dmy* on chromosome 1 is uncertain (Kasahara et al., 2007; Ichikawa et al., 2017). We intended to identify the causal gene using functional candidate gene analyses at the *r* locus then isoform differences between males and females were identified in *tetratricopeptide repeat domain 39b-like* (*ttc39bl*). Furthermore, we found that the knockout the *ttc39bl* had a white body color and was unable to complement the *r* locus.

**Figure 1.**
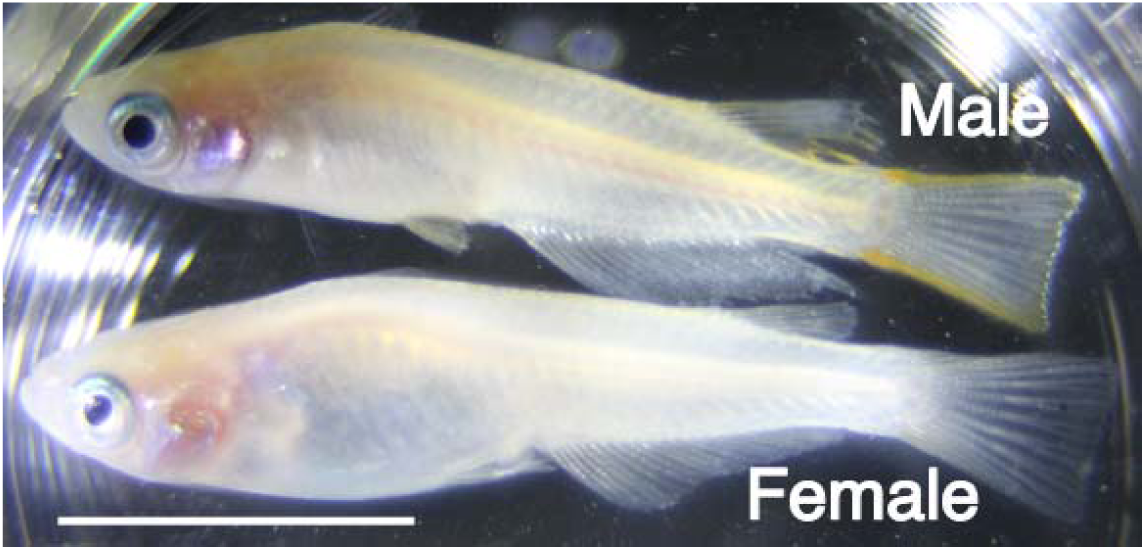
Photograph of Hd-rRII1 Adult stage Hd-rRII1. The top is male and the bottom is female Hd-rRII1. The male has orange colored xanthophores. The female, on the other hand, has uncolored xanthophores and is white. The white bar indicates 10 mm.

## Results

### Absence of the full-length *ttc39bl* mRNA in HdrR-II1 female

We scrutinized genes at the *r* locus showing differentially expressed between males and females or involvement in lipid metabolism. As a first step, we performed RNA sequencing using skins on the scales and searched for differentially expressed genes between males and females. We detected 88 up-regulated genes and 59 down-regulated genes in females, however, could not identify genes involved in lipid metabolism among the differentially expressed genes located on chromosome 1. As a next step, we took positional candidate approach then we focused on *ttc39bl* for detailed analysis because of the following reasons, even though *ttc39bl* was not differentially expressed in males and females. There are reasons to select *ttc39bl* as a candidate: *ttc39b* is known to be involved in lipid metabolism (Teslovich et al., 2010; Hsieh et al., 2016). The *ttc39b* is associated with carotenoids amount in Cichlid (Ahi et al., 2020). In clownfish, the *ttc39b* is more strongly expressed in orange stripe with many xanthophores than in white stripe with few xanthophores (Salis et al., 2019). Additionally, the *ttc39bl* is a gene near *abdg2d* (Fig.S1). We identified that HdrR-II1 had multiple *ttc39bl* mRNA isoforms and the reference isoform was only expressed in males (Fig. 2 and S2).

**Figure 2.**
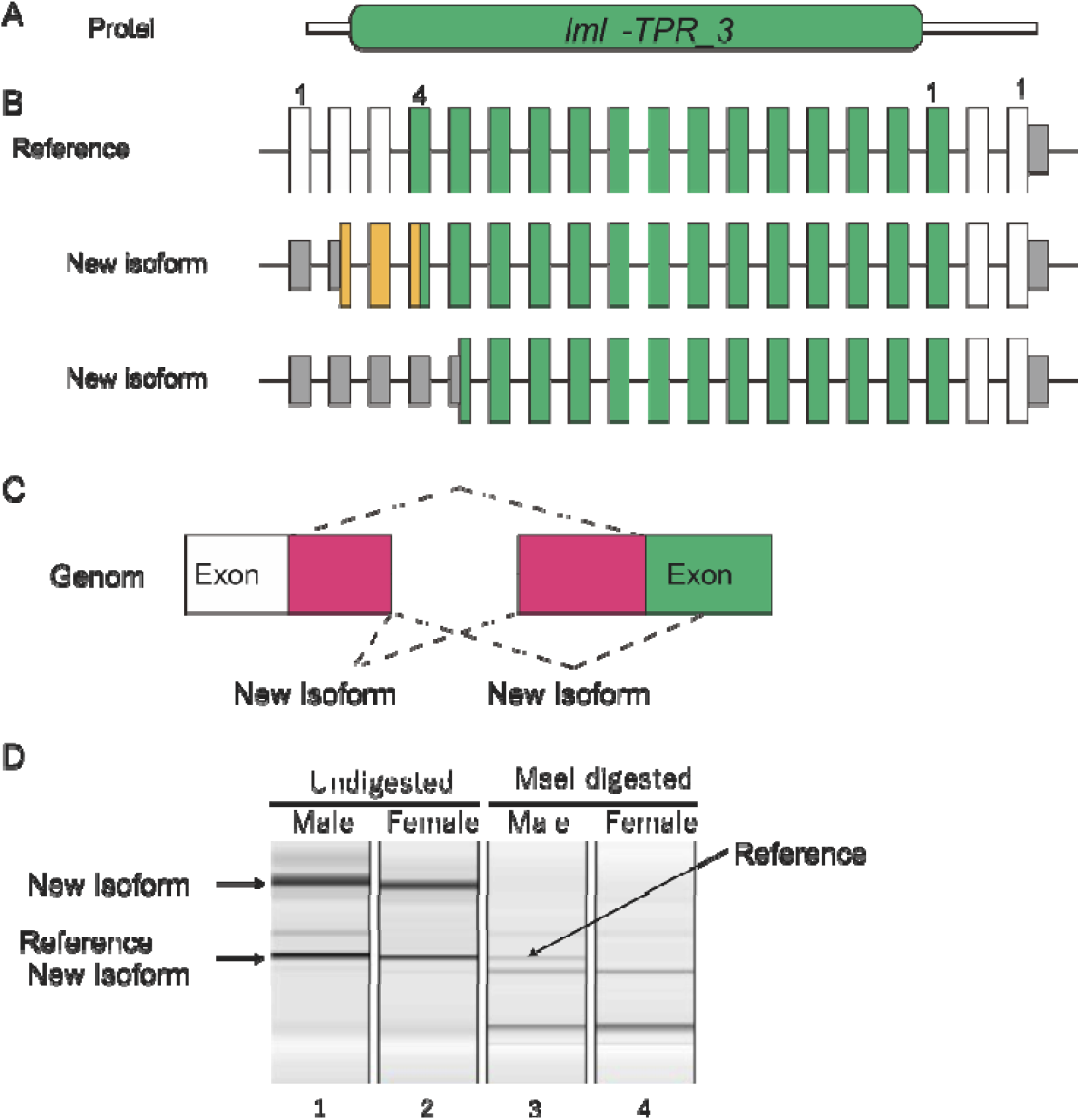
Only Hd-rRII1 male has a reference isoform of ttc39bl (A) Scheme of the domain structure of Ttc39bl. Ttc39bl is a protein consisting of 580 amino acids. It has a conserved Iml2-TPR_39 (Inclusion body clearance protein Iml2/Tetratricopeptide repeat protein 39) domain between amino acids 14 and 565 (green). (B) Scheme of exon structure of ttc39bl. The ttc39bl has 19 exons. Vertical boxes indicate exons, white boxes indicate coding regions, green boxes indicate coding regions of the Iml2-TPR_39 domain, grey boxes indicate untranslated regions, and yellow boxes indicate coding region specific in new isoform1. (C) Scheme of splicing patterns of isoforms. The white box indicates exon, the red boxes indicate coding region in new isoforms, and the green box indicates the Iml2-TPR_39 domain cording region. (D) Electrophoresis pattern of RT-PCR product and MseI digested PCR product. We designed primers ttc39bl-Ex2-L and ttc39bl-Ex5-R and performed RT-PCR using skin RNA (lanes 1 and 2). The reference isoform has no MseI site, but new isoform 1 and 2 had it. Therefore, the PCR products were digested with MseI and electrophoresed. As a result, an uncleaved band remained in the male MseI digested product (lane 3, arrow). On the other hand, all bands were shortened in the female MseI digest (lane 4).

We found at least three mRNA isoforms of the *ttc39bl* in the HdrR-II1 strain (Fig. 2 and S2). One was a reference isoform and the rest were new isoforms. The Ttc39bl domain was analyzed by InterProScan (https://www.ebi.ac.uk/interpro/), and the conserved Iml2/Tetratricopeptide repeat protein 39 (*lml2-TPR_39*) domains among TTC39A, B, and C (TTC39 domain) were found to be encoded by exon 4 to exon17 of the reference isoform (Fig. 2A). The reference isoform is the only mRNA harboring all amino acids of the TTC39 domain since frameshift occurs in new two isoforms. The *ttc39bl* mRNA has only one annotation, ENSORLT00000039395.1, in ASM223467v1 of ensemble, which corresponds to the reference isoform. Other 2 new isoforms derived from alternative splicing between exon 3 and 4 (Fig. 2B, C and S2). The sequencing inferred from the RNA-seq analysis revealed the following: The new isoform 1 encodes a protein that differs from the reference isoform by the N-terminal 28 amino acids due to alternative splicing of exons 3 and 4 (Fig. 2C and S2). The new isoform 2 encodes a protein that is 57 amino acids shorter at the N-terminus than the reference isoform due to alternative splicing of exons 3 and 4 (Fig. 2B and C). RNA sequencing suggested that only HdrR-II1 male has the reference isoform (Fig. S2). To confirm this, we performed RT-PCR using mRNA from skin, followed by MseI digestion. As the result, indeed only males had the reference isoform (Fig. 2D). Therefore, it was suggested that biological functions of *ttc39bl* might be lost in HdrR-II1 females.

### The *r* locus has 821 bp insertion in *ttc39bl*

The new isoforms 1 and 2 are produced by alternative splicing between exon 3 and exon 4, suggesting that the mutation responsible for the white body color might present in this region. Therefore, primers were designed for exons 2 and 5 and PCR was performed with genomic DNA as a template. We also used other strains of medaka, OK-Cab (orange body color) and White medaka, to examine whether this difference is the common cause of the white and orange body colors. Surprisingly, HdrR-II1 males and OK-Cab only produced a 2 kb band instead of a 2.8 kb band expected from the reference (ASM223467v1) (Fig. 3A). Only the 2 kb band was appeared in the orange medaka (HdrR-II1 male and OK-Cab), meanwhile only the 2.8 kb band was appeared in the white medaka (HdrR-II1 female and White medaka) (Fig. 3A). Since the HdrR-II1 mutation was originally derived from white medaka (Aida 1921; Yamamoto 1975; Hyodo-Taguchi 1996), these results strongly suggested that the 2.8 kb product was responsible for the white color mutation. Both the 2.8 kb and 2 kb bands were sequenced by Sanger method. The 2.8 kb band showed exact match to the reference sequence and the 821 nucleotides in intron 3 was lacking in the 2 kb band sequence of the orange medaka (Fig. 3B). This suggested that the 821-base insertion disrupted the orange medaka exon 3 of ttc39bl, splitting it into exons 3 and 4 of the reference sequence, causing de novo splicing to occur and causing loss of ttc39bl function (Fig. 3B). Also, the reference sequence is probably derived from the X chromosome of HdrR-II1.

**Figure 3.**
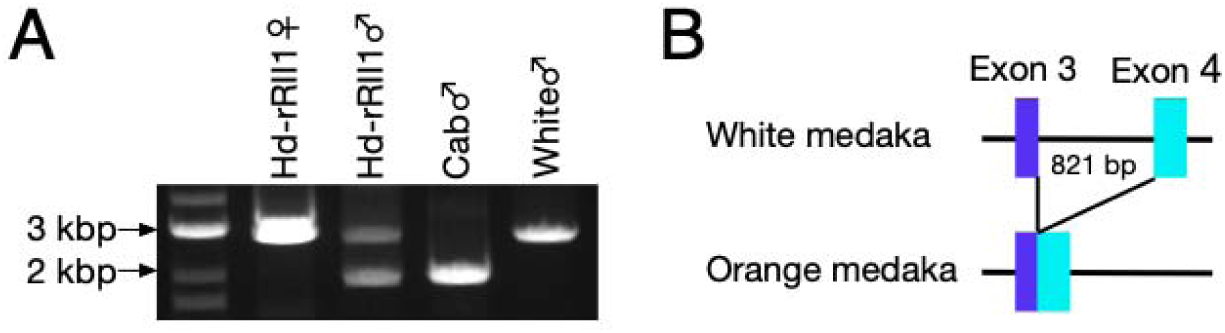
White medakas have 821 base pair insertion, but orange medakas do not. (A) Genomic PCR products of ttc39bl. White body color medakas (white male and Hd-rRII1 female) have only a 3 kb band. On the other hand, orange body color medakas (Hd-rRII1 male and Cab male) have 2 kb and 3 kb bands. (B) Schema of a causal mutation in ttc39bl. Sanger sequencing elucidated that the reference sequence of ttc39bl was derived from Hd-rRII1 and exon 3 and 4 are one exon in the wild-type genome. In other words, the intron 3 of reference sequence is the causal mutation at the r locus, disrupting exon 3 of wild-type ttc39bl.

### The *ttc39bl* knockout by CRISPR/Cas9 cannot complement the *r* locus

In order to confirm whether non-functional *ttc39bl* indeed results in the white body color, we introduced a stop codon at exon 3 (exon 4 in the reference genome) of the KO-Cab genome by using CRISPR/Cas9. We designed a guide RNA (gRNA) that introduces a double strand break in the TTC39 domain (Fig. 4A). One of the injected CRISPANT showed white color (Fig. 4B). We raised this CRISPANT and crossed it with a male HdrR-II1 and found white females in the offsprings (Fig. 4C). Sanger sequencing confirmed the 28 nucleotides deletion in white offsprings (Fig. 4D). This deletion introduced a frameshift and a new stop codon in *lml2-TPR_39* domain. Surprisingly, the other white fish had a 3 bases deletion mutation losing the 41st asparagine. This mutation results in the loss of only one amino acid in the *lml2-TPR_39* domain but failed to complement the *r* allele. This suggests that the complete presence of the *lml2-TPR_39* domain is required for xanthophores to accumulate carotenoid pigments. We thought that the causative mutation at the *r* locus was a splicing defect, but it was caused by an exon disruption by the 821 bases insertion. Taken together, we concluded that *ttc39bl* is the responsible gene for the *r* locus.

**Figure 4.**
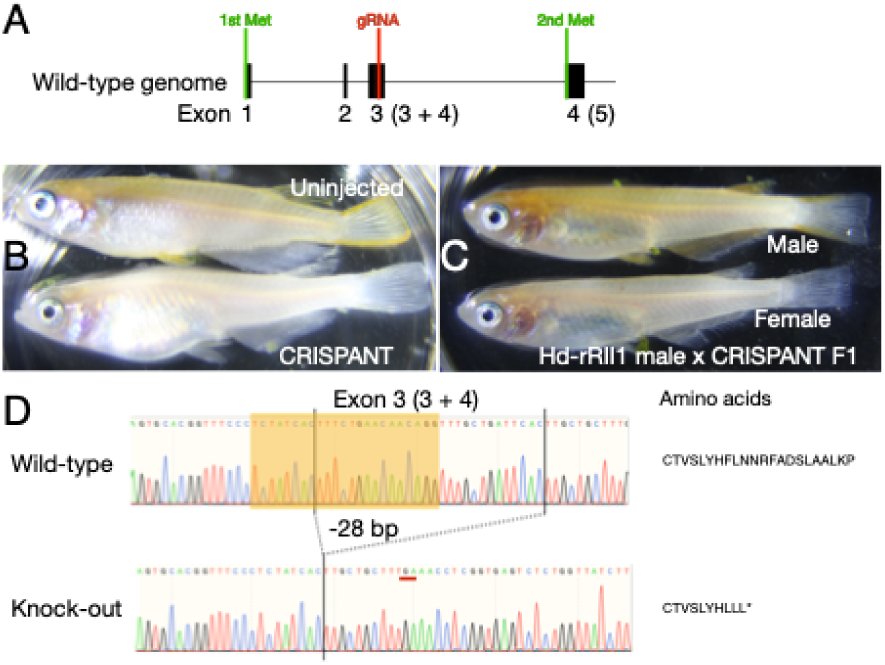
ttc39bl CRISPANT and knock-out have white body color. (A) Position of guide RNA (gRNA) (red line) and methionine codons (green line). Numbers in parentheses are exon numbers in reference genome sequence. (B) ttc39bl CRISPANT photograph. The ttc39bl CRISPANT has white body color. (C) F1 photograph derived from a cross between Hd-rRII1 male and ttc39bl CRISPANT female. Fish heterozygous for the ttc39bl mutant and the r locus had white body color. Thus, the ttc39bl mutation does not complement the r locus. (D) Sanger sequencing showed that the ttc39bl knock-out has 28 base pair deletion in exon 3. Deletion induced frameshift and early stop codon. Orange-box indicates gRNA target region and PAM sequence. The red line indicates a stop codon induced by the deletion. Numbers in parentheses are exon numbers in reference genome sequence.

## Discussion

White medaka had been bred and maintained for more than 200 years with himedaka because they were drawn in pictures at the Edo period. It is very impressive that medaka, which has no industrial value, has been loved for such a long period. In this study, we tried to identify the responsible gene for the *r* locus resulting in the white color, which had been unknown for more than 100 years.

In the current study, we demonstrated that the full-length isoform of *ttc39bl* was lost in HdrR-II1 female. We also found that the exon 3 of *ttc39bl* is disrupted due to 821 bases insertion in white medaka but not in orange medaka. Finally, we show the *ttc39bl* mutation induced by CRISPR/Cas9 could not complement the *r* locus. Taken together, our results indicate that the responsible gene for the *r* locus would be *ttc39bl*. Our results provide direct evidence that the ttc39b gene is an essential gene for carotenoid collar accumulation.

Ttc39b is expected to be involved in carotenoid accumulation through lipid metabolism. *TTC39B* is associated with serum concentrations of high-density lipoprotein cholesterol in human (Teslovich et al., 2010). The defect of *Ttc39b* inhibits dietary cholesterol absorption and accumulation in mice (Hsieh et al., 2016). In *Ttc39b* knock-out mice, the loss of Ttc39b suppresses liver X receptor (LXR) ubiquitination and increases the stability of LXR protein (Hsieh et al., 2016). In enterocytes, LXR protein increases the expression of *ATP-binding cassette transporter A1* (*Abca1*) and promotes high-density lipoprotein cholesterol production. Since carotenoids are lipophilic, their amounts vary depending on cholesterol level. *TTC39B* is highly expressed in xanthophore/erythrophore-rich regions suggesting that it is also required for these cells to take up carotenoids. Presumably, *TTC39B* is involved in the process of absorption and accumulation of food-derived carotenoids.

Medaka has two *TTC39B* orthologues, *ttc39b* (ENSORLG00000027310) on chromosome 18 and *ttc39bl* on chromosome 1. Since white medaka is healthy and fertile, probably *ttc39b* complements the function of *ttc39bl*. Future studies are needed to understand the function of the *ttc39bl* outside of body color.

The 821 bp insertion in *ttc39bl* of white medaka was found in the genomes of the HdrR-II, the HNI, and HSOK lines with identities of 52, 65, and 16 % (length > 800, and E-value is 0), respectively. However, an identical sequence of the insert was only found in *Oryzias latipes* and not in other *Oryzias* species, suggesting that the origin of insertion was relatively new. Curiously, the homologous sequence exists in the genome of *Acanthopagrus latus* (85% identities (410/483), and E-value is 3 x 10^-126^) and *Salarias fasciatus* (83% identities (220/266), and E-value is 1 x 10^-50^), which are not a close relative of medaka. The insertion does not appear to encode any protein as a stop codon appears immediately in any reading frame. Future research will elucidate the origin of the insertion.

In some sexually dimorphic fishes, genes responsible for sex-dependent body color differences have been reported in the sex-determining genomic region (Kottler and Schartl 2018). The sex chromosome of medaka contains *ttc39bl*, which is identified in this study, and *slc2a15b*, which is involves in the pigmentation of leucophores. The TTC39B highlighted in this study has also been reported to be in the sex chromosome in birds (Hooper et al., 2019). However, although the distribution of yellow pigment cells is slightly different between male and female medaka, there is no difference in body color. In females, the xanthophores are uniformly distributed, but in males they are concentrated at the edges of the fins (Hama 1975; Iwamatsu 1997). During the breeding season in males, melanophores and xanthophores become darker in color, and leucophroes develop at the tips of fins and the snout to the eyes. It has been reported that the nuptial coloration of medaka is lost in castrated males and induced in females by testosterone treatment, suggesting stronger hormonal control than genetic control (Iwamatsu 1997).

This study showed that *ttc39bl*, one of *TTC39B* orthologue, is indispensable for carotenoid accumulation of xanthophores. However, the interactions of *ttc39bl* with other genes involved in carotenoids accumulation such as *scarb1* and *plin6* are unclear. It is also unclear whether *ttc39b* is responsible for the diversity of yellow to orange body color found in medaka. It is necessary to further elucidate the genes involved in the accumulation of carotenoids and to identify genetic polymorphisms involved in body color diversity in the future.

## Experimental procedures

### Animals

The HdrR-II1 (Strain ID: IB178) and OK-Cab (Strain ID: MT830) strains used in this study were obtained from the National BioResource Project (NBRP), Medaka (https://shigen.nig.ac.jp/medaka/). White medaka was obtained from a pet shop. All three medakas have a mutation in *solute carrier family 45 member 2* (*slc45a2*) and the black pigment of the skin is lost. All medakas were reared at 26.0L on 14-hr light/10-hr dark cycle.

### The gene list at the *r* locus

The *r* locus has been reported to be at 0.18 cM telomeric from 51H.7F, the flanking marker of *dmy* (Matsuda et al., 2002). Location of the 51H7.F was searched by BLAT search for medaka 2005 genome (oryLat2) in UCSC Genome Browser (http://genome.ucsc.edu) (Kent 2002) and for medaka 2017 genome (ASM223467v1) in Ensembl genome browser (https://ensembl.org) (Howe et. al., 2021). In both genome sequences, the 51H7.F is locating on *ATP-binding cassette sub-family G member 2d* (*abcg2d*) (ENSORLG00000006979) (Fig 2). Synteny around *abcg2d* among medaka, pufferfish, and green spotted pufferfish were investigated using Genomicus synteny browser (https://www.genomicus.bio.ens.psl.eu/) (Louis et al. 2015). Then, we listed the genes between *abcg2d* and *lef1* in the 2017 genome. Finally, those two were merged to create the genes list of the *r* locus.

### RNA extraction and RNA sequencing analysis

Medaka skin was collected by stripped off scales with scalpel and subjected to RNA preparation. Total RNAs were extracted using TRIzol (Ambion) and purified using RNeasy mini Kit (QIAGEN) according to the manufacturer’s protocols. Samples were sequenced on MGI DNGSEQ-G400 150 bp paired end.

Removal of adapter sequences and low-quality reads were performed using Trimmomatic (v.0.39) (Bolger et al., 2014). The filtered reads were mapped to Japanese medaka HdrR (ASM223467v1) and HNI (ASM22347v1) genome using STAR (v2.7.10a) (Dobin et al., 2013). Quantification of genes and transcripts was performed using featureCounts (v.2.0.1) (Liao et. al., 2019). Differential expression analysis was performed using edgR (v.3.36.0) and limma (v.3.50.1) (Law et. al., 2016). Transcript isoforms were examined by Integrative Genome Viewer (v.2.8.0) (Robinson et al., 2011).

### Protein domain search

We performed the domain search of ttc39bl isoforms by InterProScan (https://www.ebi.ac.uk/interpro/) (Jones et. al., 2014).

### RT-PCR and MesI digestion

RNeasy (QIAGEN) and ReverTra Ace (Toyobo) were used to prepare cDNA according to the manufacturer’s protocol. The primers used for amplification are shown in Table S1. The PCR conditions were as follows; one cycle at 98L for 2 min, 35 cycles at 98L for 10 sec, 54L for 10 sec, and 68L for 30 sec followed by 68L for 1 min. The PCR products and MseI digested samples were electrophoresed using a Standard Cartridge Kit on Qsep 100 DNA Analyzer (BiOptic Inc).

### CRISPR/Cas9 gene targeting and mutation identification

The sgRNA were designed using CRISPRdirect (https://crispr.dbcls.jp) (Naito et. al., 2015). The sgRNA and CRISPR/Cas9 protein were purchased from INTGRATED DNA TECHNOLOGIES. To generate Cas9 ribonucleoprotein complex, we mixed Cas9 protein (5 µM), sgRNA (5 µM), KCl (300 mM), Tris-HCl (pH 8.0) (10 mM), Phenol Red (0.025 %) in 10 µL (Wu et al., 2018). The mixture was incubated at 37L for 5 minutes. The sgRNA sequence is ‘TCTATCACTTTCTGAACAAC’.

Microinjection was performed using a needle made by pulling the glass capillary G-1 (NARISHIGE) with PC-10 (NARISHIGE) on a manipulator and applying pressure with a 50 mL syringe (TERUMO).

Genomic DNA was purified from fin clips with HotSHOT method (Truett et al. 2000). The target entire region of *ttc39bl* was amplified with Tks Gflex DNA Polymerase (Takara Bio). The PCR condition was as follows; one cycle at 98L for 2 min, 35 cycles at 98L for 10 sec, 55L for 15 sec, and 68L for 20 sec followed by 68L for 40 sec. The PCR products were treated with ExoSAP-IT PCR Product Cleanup (ThermoFisher) according to the manufacturer’s protocol and directly sequenced by Sanger method.

### Genomic sequence

Genomic DNAs were extracted from eyeballs using smart DNA prep (m) (Analytik Jena GmbH) according to the manufacturer’s protocol. The PCR condition was as follows; one cycle at 98L for 2 min, 35 cycles at 98L for 10 sec, 55L for 15 sec, and 68L for 2 min 30 sec followed by 68L for 5 min with Tks Gflex DNA Polymerase (Takara Bio). The PCR products were cloned into pCR Blunt II-TOPO vector using Zero Blunt TOPO PCR Cloning Kit (ThermoFisher) according to the manufacturer’s protocol. Colonies were picked up into PCR mixture and amplified. The PCR products were treated with ExoSAP-IT PCR Product Cleanup (ThermoFisher) and sequenced directly with Applied Biosystems 3500 Genetic Analyzer. The primers used for amplification and sequencing are shown in Table S1. The sequences were analyzed using by BLAT search using UCSC Genome Browser.

### Primer information

RT-PCR was performed using ttc39bl-Ex2 (AGTGGACGTTTTTGAAGATG) and ttc39bl-Ex5 (TAGCCCACTGCATGGTAC). Genomic PCR in Figure 3 was performed using ttc39bl-Ex3 (CAGCACAAATTGACCTGG) and ttc39bl-Ex5. Genomic sequencing of white and orange body medaka was performed using ttc39bl-Ex3 and ttc39bl-intron4 (GCGCAAATCTAAACTGCACAC). Cas9-induced mutations detection sequence was performed using ttc39bl-Ex3 and ttc39bl-intron4.

## Acknowledgments

We are grateful to NBRP Medaka (https://shigen.nig.ac.jp/medaka/) for providing HdrR-II1 (Strain ID: IB178) and OK-Cab (Strain ID: MT830). This work was supported by the Japanese Agency for Medical Research and Development under grant number 22ek0109484h0003.

## Author contributions

TK designed research, performed research and prepared all figures; TK and II wrote the manuscript.

