## Supplemental Figures for "*ttc39bl,* one of the two duplicated paralogs of the *tetratricopeptide repeat domain 39B* gene, is essential for carotenoid coloration in medaka (Oryzias latipes) and is the gene responsible for the *r* locus"

**Supplementary material**

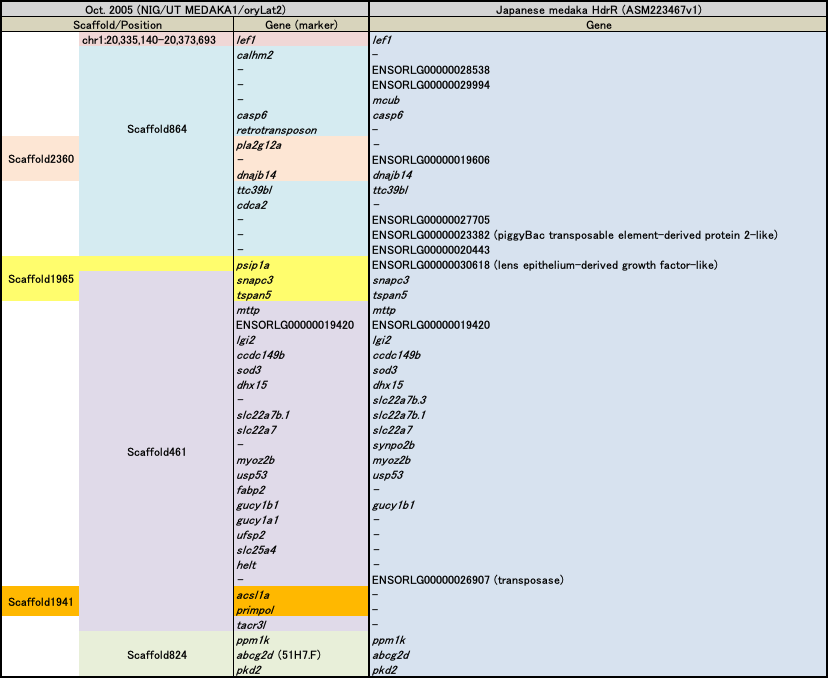

Figure S1 Gene list at the r locus

Two major genome sequences of medaka were registered, oryLat2 (in UCSC Genome Browser) and ASM223467v1 (in Ensembl). Colors in the oryLat2 column show each scaffold. For example, *pla2g12a* is in scaffold864 and scaffold2360. Hyphen indicates no corresponding gene in the genome.

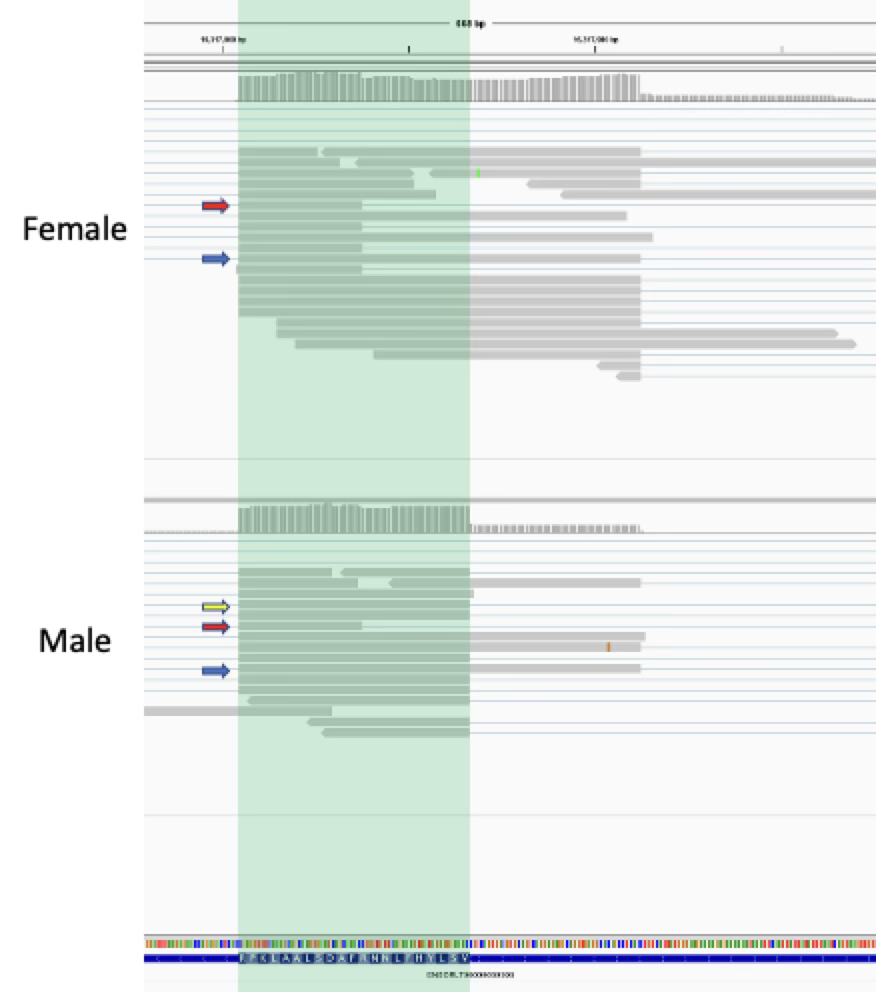
Figure S2 Exon 4 of ttc39bl mapping results

The *ttc39bl* exon 4 mapping result of RNA sequencing. The green highlight indicates the region corresponding to exon 4 of the reference sequence. The yellow arrow indicates the read corresponding to the reference isoform. The red arrows indicate the read corresponding to new isoform 1. The blue arrows indicate the read corresponding to new isoform 2. The female has no reference isoform.
